# Interleukin-1 receptor antagonist treatment in acute ischaemic stroke does not alter systemic markers of anti-microbial defence

**DOI:** 10.1101/587881

**Authors:** Laura McCulloch, Stuart M. Allan, Craig J. Smith, Barry W. McColl

**Affiliations:** UK Dementia Research Institute, University of Edinburgh, Edinburgh, United Kingdom; Division of Neuroscience and Experimental Psychology, School of Biological Sciences, Faculty of Biology, Medicine and Health, University of Manchester, Manchester Academic Health Science Centre, Manchester, UK.; Division of Cardiovascular Sciences, University of Manchester, Manchester, UK and Greater Manchester Comprehensive Stroke Centre, Manchester Centre for Clinical Neurosciences, Manchester Academic Health Science Centre, Salford Royal NHS Foundation Trust, Salford, UK

**Keywords:** stroke, IL-1Ra, antibodies, complement, infection

## Abstract

**Aim:** Blockade of the cytokine interleukin-1 (IL-1) with IL-1 receptor antagonist (IL-1Ra) is a candidate treatment for stroke entering phase II/III trials, which acts by inhibiting harmful inflammatory responses. Infection is a common complication after stroke that significantly worsens outcome and is related to stroke-induced deficits in systemic immune function thought to be mediated by the sympathetic nervous system. Therefore, immunomodulatory treatments for stroke, such as IL-1Ra, carry a risk of aggravating stroke-associated infection. Our primary objective was to determine if factors associated with antibody-mediated antibacterial defences were further compromised in patients treated with IL-1Ra after stroke.

**Methods:** We assessed plasma concentrations of immunoglobulin isotypes and complement components in stroke patients treated with IL-1Ra or placebo and untreated non-stroke controls using multiplex protein assays. Activation of the SNS was determined by measuring noradrenaline, a major SNS mediator.

**Results:** There were significantly lower plasma concentrations of IgM, IgA, IgG1 and IgG4 in stroke-patients compared to non-stroke controls, however there were no differences between stroke patients treated with placebo or IL-1Ra. Concentrations of complement components associated with the classical pathway were increased and those associated with the alternative pathways decreased in stroke patients, neither being affected by treatment with IL-1Ra. Noradrenaline concentrations were increased after stroke in both placebo and IL-1Ra-treated stroke patients compared to non-stroke controls.

**Conclusion:** These data show treatment with IL-1Ra after stroke does not alter circulating immunoglobulin and complement concentrations, and is therefore unlikely to further aggravate stroke-associated infection susceptibility through reduced availability of these key anti-microbial mediators.

## INTRODUCTION

Blocking the actions of the inflammatory cytokine interleukin-1 (IL-1) using a highly selective IL-1 receptor antagonist (IL-1Ra) reduced injury and improved outcome in multiple experimental animal models of cerebral ischemia and is in ongoing clinical stroke trials ^1–4^. The inflammatory-modifying properties of IL-1Ra may confer protective effects to the brain after stroke, however due to its potential for immunosuppression, it may also compromise systemic immune responses important for defence against infection. Systemic immune dysregulation is particularly important to consider in the context of stroke as patients are highly susceptible to infection which likely involves roles for stroke-induced impairments in some immune functions ^5^.

We have previously shown deficits in early antibody responses, particularly IgM, associated with innate-like B cells in both experimental animals and stroke patients that may contribute to post-stroke infection susceptibility ^6^. Il-1β is reported to induce IgM production in innate-like B cells ^7^, therefore treatment with IL-1Ra may inhibit these important anti-microbial effects. We assessed if markers associated with antibody-mediated antibacterial defences were compromised in patients treated with IL-1Ra after stroke. Plasma IgM, IgG1, IgG4 and IgA immunoglobulin concentrations were reduced after stroke and this was not further altered by treatment with IL-1Ra. Assessment of complement components indicated induction of the classical pathway of complement activation after stroke but inhibition of the alternative pathway without modulation by IL-1Ra. Plasma noradrenaline was increased after stroke and also not influenced by treatment with IL-1Ra. In summary, our data suggest treatment with IL-1Ra is unlikely to aggravate antibody-associated immune function deficits induced by stroke.

## METHODS

### Participants and study procedures

In brief, patients ≥ 18 years of age with a clinical diagnosis of stroke within 6 h of stroke onset were eligible. Exclusion criteria included National Institutes of Health Stroke Scale (NIHSS) score of ≤ 4, pre-stroke modified Rankin Scale (mRS) score of ≥4 or rapidly improving neurological deficit. Patients were randomly assigned to treatment with recombinant methionylated human IL-1Ra (n=17) or placebo (n=17) stratified by age (<70 and ≥ 70 years), baseline stroke severity (NIHSS score 4-9, 10-20, ≥ 21) and time since stroke onset (<4 or ≥ 4 h) but not by sex. IL-1Ra was initially administered as an IV loading dose of 100 mg over 60 seconds followed by 72 h of consecutive infusions at 2 mg/kg/h. Full patient baseline characteristics and stratification of groups are provided in Supplementary Table e-1.

Non-stroke control patients (n=13) of a similar age range with no previous history of stroke or transient ischemic attack were also recruited. Control patients were living independently at home, free of infection and able to provide written, informed consent. Controls were matched to stroke patients (6 to patients receiving IL-1Ra and 7 to patients receiving placebo) on a basis of age (±5 years), sex and degree of atherosclerosis.

### Blood sampling

Venous blood samples were collected prior to initiation of treatment (admission), at the next 9am time point (if admission was before 7 am or after 11am), and then at 9 am at 24 h, 2 d, 3 d, 4 d and at 5-7 d after stroke, into tubes containing a final concentration of 10 μg/ml pyrogen-free heparin and wrapped in cool packs. Control patients were sampled at 9 am and also at matched patient admission time (2 h) if this was not between 7 and 11 am. Samples were centrifuged 1 h after collection at 2000 *xg* for 30 min at 4°C. Plasma was separated and frozen in aliquots at −70 °C until further analysis.

### Standard Protocol Approvals, Registrations, and Patient Consents

This study involved tertiary analysis of plasma samples taken from a randomised, placebo-controlled phase II trial originally designed to determine the safety and biological activity of intravenous (IV) IL-1Ra ^4^. The online clinical trials registries ClinicalTrials.gov and ISRCTN went live online during the year 2000, at which time online trial registration was a relatively new recommendation. The original IV IL-1Ra trial was set-up in 2000, and commenced Feb 2001 and therefore this trial was not officially registered. Ethical approval for reanalysis of the samples was obtained through the Health Research Authority National Research and Ethics Service Committee (16/NW/0853).

### Luminex analysis of immunoglobulins and complement components

Immunoglobulins and complement components were measured in plasma samples using MILLIPLEX^®^ multiplex assays. Patient details were blinded from samples and coded samples were randomised across plates for analysis. The MILLIPLEX^®^_MAP_ Human Isotyping Magnetic Bead Panel-Isotyping Multiplex Assay (HGAMMAG-301K-06, Merck Millipore Corporation, Billerica, MA, USA) was used to measure IgG1, IgG2, IgG3, IgG4, IgA and IgM. MILLIPLEX^®^_MAP_ Human Complement Panel 1 was used to measure C2, C4b, C5, C9, Mannose-binding lectin (MBL), Factor D (Adipsin) and Factor I (HCMP1MAG-19K, Merck Millipore Corporation). Many samples had concentrations of Factor D and Factor I below the detection range of the standard curve and so results for these analytes are not reported. MILLIPLEX^®^_MAP_ Human Complement Panel 2 was used to measure C1q, C3, C3b/ iC3b, C4, Factor B, Properdin and Factor H (HCMP2MAG-19K, Merck Millipore Corporation). Samples were assayed as singlets and all samples, standards and quality controls were prepared in accordance with the manufacturer’s instructions. Samples were incubated with beads on plate for 1 h (Isotyping assay) or overnight (Complement assays) at 4°C and washes carried out using a magnetic plate washer. Plates were analysed using a Magpix™ Luminex^®^ machine and Luminex xPonent^®^ software.

### Measurement of Noradrenaline

Noradrenaline was measured in plasma samples using a Noradrenaline ELISA kit (BA E-5200; LDN^®^, Nordhorn, Germany). Patient details were blinded from samples and coded samples were randomised across plates for analysis. Samples were assayed as singlets and all samples, standards and quality controls were prepared in accordance with the manufacturer’s instructions where noradrenaline is extracted from plasma using a cis-diol-specific affinity gel, acylated, enzymatically converted and then measured by ELISA. Optical density at 450 nm was measured using an MRX microplate Reader (Dynatech Labs, Chantilly, VA).

### Statistical analyses

All immunoglobulin and complement components were measured in μg/ml and the D’Agostino and Pearson omnibus test was used to determine Gaussian distribution of sample data. As data were non-normally distributed, sample values were log_10_-transformed. As the precise kinetics of individual patient responses may vary, the maximal and minimal concentrations of each mediator in the first 7 d after stroke were compared to non-stroke controls. Maximal and minimal concentrations from IL-1Ra-treated and placebo-treated stroke patients and non-stroke controls were compared by one-way ANOVA with Bonferonni correction. Noradrenaline concentrations were measured in ng/ml and the D’Agostino and Pearson omnibus test was used to confirm Gaussian distribution of sample data. Maximal and minimal noradrenaline concentration from IL-1Ra-treated and placebo-treated stroke patients and non-stroke controls were compared by one-way ANOVA with Bonferonni correction. Data analysis was performed using GraphPad Prism 6.0 statistical analysis software and for all experiments, values of *P* ≤ 0.05 were accepted as statistically significant.

### Data availability

Anonymised data will be shared on reasonable request from any qualified investigator

## RESULTS

### Plasma IgM concentration is reduced after stroke and is not affected by treatment with IL-1Ra

Immunoglobulin M (IgM) is the predominant immunoglobulin isotype associated with early B cell antibody responses to infection by innate-like B cells which we have previously shown to be depleted after experimental stroke in mice ^6, 8, 9^. Lower minimum concentrations of IgM were measured after stroke in comparison to non-stroke controls, and no difference was found between placebo and IL-1Ra treated patients. (Figure 1A). Maximum IgM concentrations in the first 7 days after stroke were also assessed and did not significantly differ in IL-1Ra or placebo treated patients in comparison to non-stroke controls (**Figure e-1A**). This indicates that the reduced minimum IgM concentration measured over the first 7d reflects an actual reduction in circulating IgM in stroke patients and is not an artefact of increased variance in IgM concentration after stroke.

**Figure 1.**
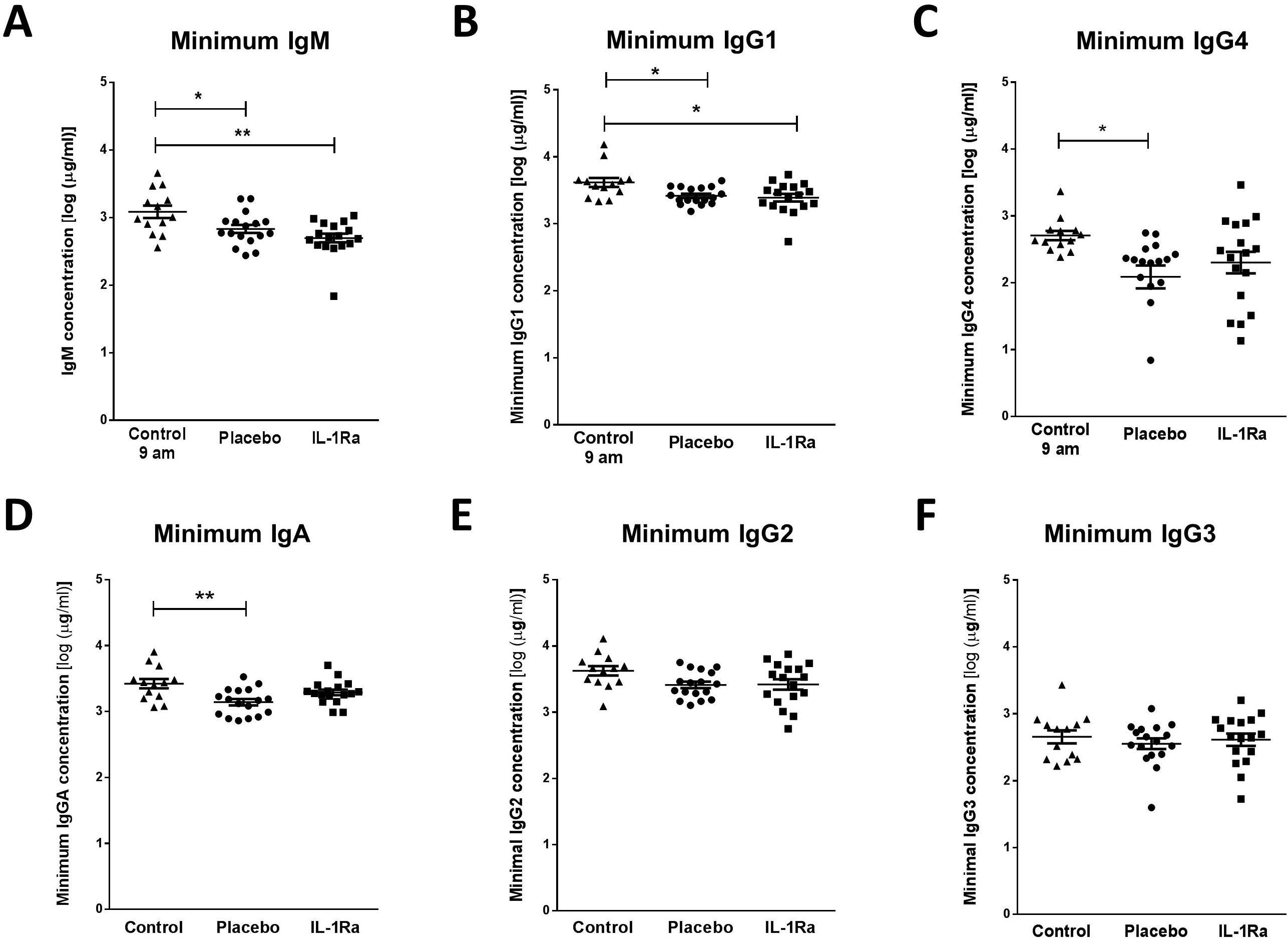
Reduced plasma IgM, IgA, IgG1 and IgG4 after stroke is not affected by IL-1Ra. **(A)** Minimum IgM concentration measured in the first 7 d after stroke was lower in both placebo and IL-1Ra treated patients in comparison to healthy controls. Data show mean ±SD, * *P<0.05*, ** *P<0.01*, one-way ANOVA with Bonferonni correction. **(B)** Minimum concentration of IgG1 and measured in the first 7 d after stroke was reduced in both placebo and IL-1Ra treated patients in comparison to healthy controls. Minimum IgG4 **(C)** and IgA **(D)** concentrations were reduced in placebo-treated stroke patients in comparison to healthy controls. There was no significant difference between placebo-treated and IL-1Ra-treated stroke patients. No significant difference in IgG2 **(E)** and IgG3 **(F)** concentration was detected between placebo-treated and IL-1Ra-treated stroke patients in comparison to healthy controls. Data show mean ±SD, * *P<0.05*; ** *P<0.01;* one-way ANOVA with Bonferonni correction.

### Plasma IgA, IgG1 and IgG4 concentrations are reduced after stroke and are not affected by treatment with IL-1Ra

Minimum IgG1 concentration was significantly reduced in both placebo-treated and IL-1Ra-treated stroke patients in comparison to non-stroke controls (Figure 1B). Minimum IgG4 (Figure 1C) and IgA (Figure 1D) concentrations were significantly reduced in placebo-treated stroke patients only. However, there was no significant difference in these immunoglobulins between placebo-treated and IL-1Ra-treated patients. Minimum concentrations of IgG2 (Figure 1E) and IgG3 (Figure 1F) were not significantly altered in IL-1Ra or placebo treated patients in comparison to non-stroke controls. Maximal circulating concentrations of all immunoglobulin isotypes measured in the first 7 days after stroke were also compared to non-stroke controls and no significant differences were measured in any immunoglobulin isotypes (**Figure e-1**).

### Concentrations of complement components are differentially affected by stroke and not affected by treatment with IL-1Ra

As complement components are directly associated with the antibacterial functions of immunoglobulins, we investigated stroke-induced changes in circulating complement components and if any changes observed were further influenced by treatment with IL-1Ra. Stroke induced a significant reduction in the minimum concentrations of C3b/ iC3b (Figure 2A), C3 (Figure2B), C4 (Figure 2C), Factor H (Figure 2D) and Properdin (Figure 2E) measured in the first 7 days after stroke in both placebo and IL-1Ra treated patients in comparison to non-stroke controls. Maximum circulating concentrations of these complement components measured in the first 7 days after stroke were also compared to non-stroke controls and no significant differences were seen (**Fig e-2A-E**).

**Figure 2.**
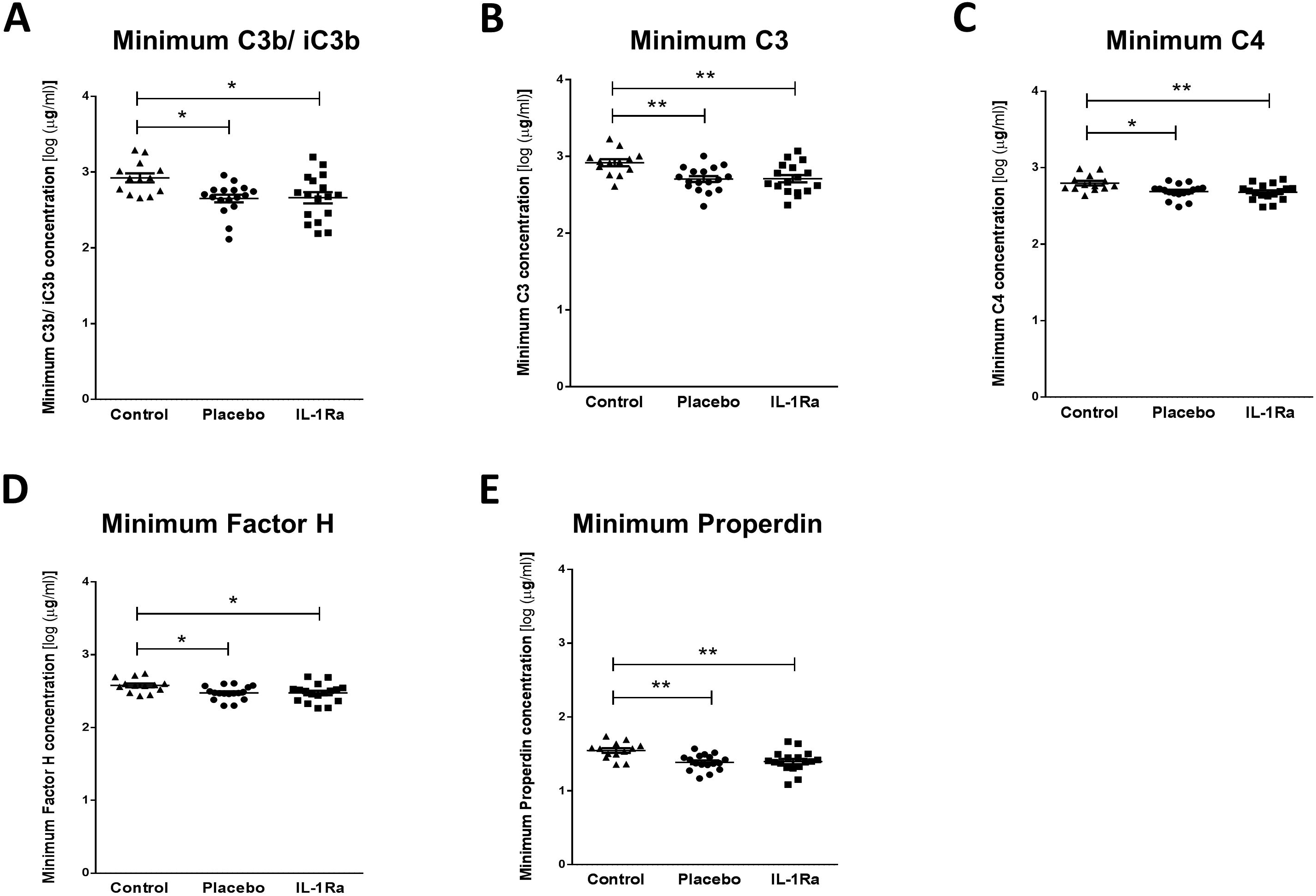
Treatment with IL-1Ra has no effect on complement components downregulated after stroke. Minimum concentrations of **(A)** C3b/ iC3b, **(B)** C3, **(C)** C4, (**D)** Factor H and **(E)** Properdin were measured in the first 7 d after stroke were reduced in both placebo and IL-1Ra treated patients in comparison to healthy controls. Data show mean ±SD, * *P<0.05*; ** *P<0.01;* one-way ANOVA with Bonferonni correction.

In contrast, stroke induced a significant increase in maximal circulating concentrations of C1q (Figure 3A), C5 (Figure 3D) and C9 (Figure 3E) in both IL-1Ra and placebo treated patients measured in the first 7 days after stroke in comparison to non-stroke controls. Maximum concentrations of C2 (Figure 3B) and C4b (Figure 3C) were increased in placebo-treated patients only however no significant difference was apparent between placebo treated and IL-1Ra treated patients for these factors suggesting IL-1Ra treatment exerts no effects additional to stroke. Minimum concentrations of these complement components measured in the first week after stroke were also compared to non-stroke controls and no significant differences were seen (Figure e-3A-E).

**Figure 3.**
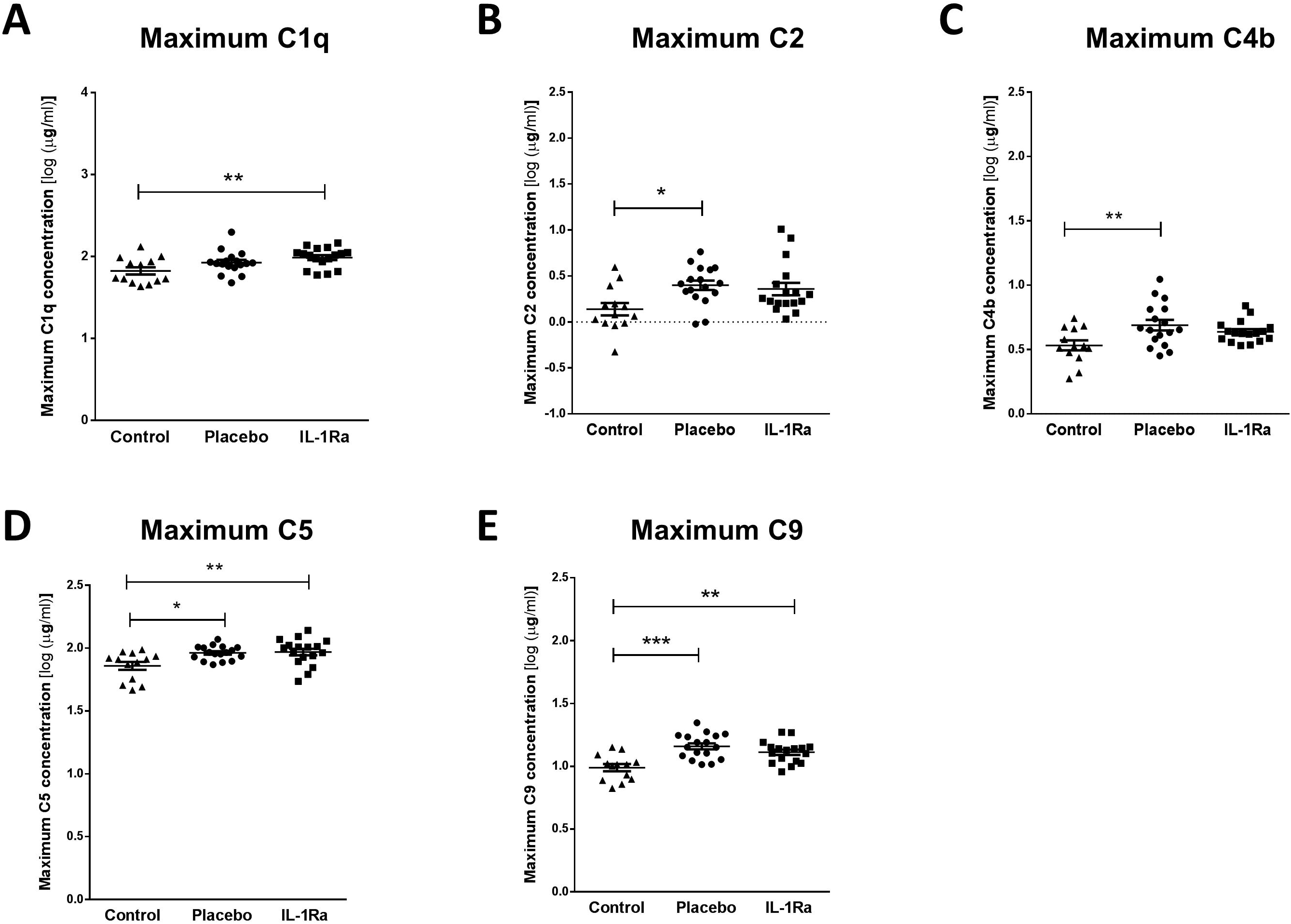
Treatment with IL-1Ra has no effect on complement components upregulated after stroke. Maximum concentrations of **(A)** C1q, **(B)** C2, **(C)** C4b, **(D)** C5 and **(E)** C9 were measured in the first 7 d after stroke were increased in both placebo and IL-1Ra treated patients in comparison to healthy controls. Data show mean ±SD, * *P<0.05*; ** *P<0.01;* one-way ANOVA with Bonferonni correction.

Minimal and maximal levels of factor B, mannose-binding lectin (MBL) and C5a measured in the first week after stroke were also compared to non-stroke controls. Concentrations of Factor B (Figure 4A, B) and MBL (Figure 4C, D) were not significantly altered by stroke or by treatment with IL-1-Ra.

**Figure 4.**
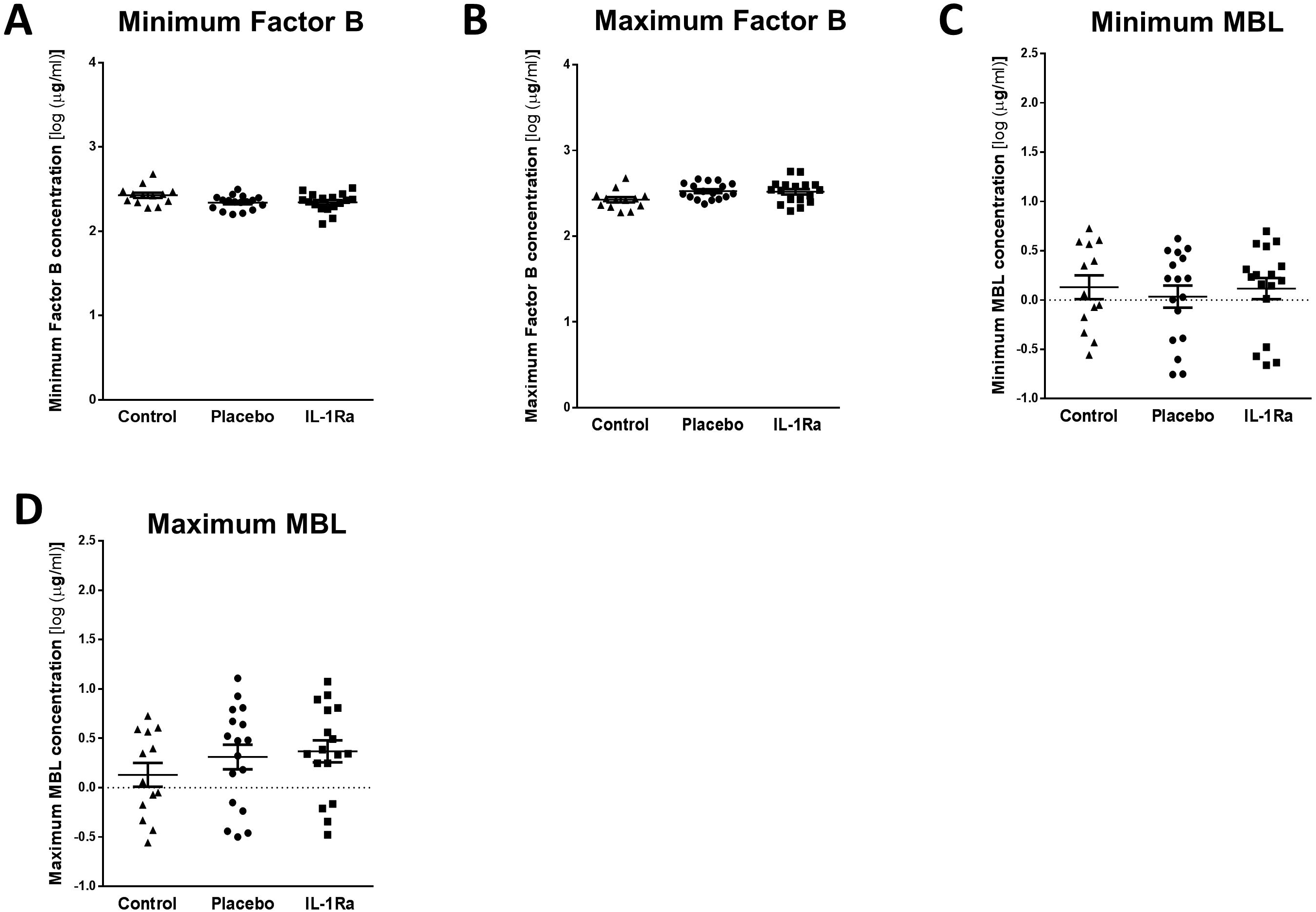
Treatment with IL-1Ra has no additional effect on complement components unaffected by stroke. Minimal and maximum concentrations of **(A, B)** Factor B and **(C, D)** MBL were measured in the first 7 d after stroke were unchanged in both placebo and IL-1Ra treated patients in comparison to healthy controls. Data show mean ±SD, one-way ANOVA with Bonferonni correction.

### Plasma noradrenaline concentration is increased after stroke and is not affected by treatment with IL-1Ra

Splenic noradrenaline levels are increased after experimental stroke and may be toxic to IgM producing B cells ^6^. Maximum noradrenaline concentration measured in the first 7 days after stroke was increased in both placebo and IL-1Ra treated patients in comparison to non-stroke controls (Figure 5A). Treatment with IL-1Ra had no additional effect on noradrenaline concentration when compared to placebo. Minimum noradrenaline concentration measured in the first 7 days after stroke was also measured and was not significantly different to non-stroke controls (**Figure e-4**), or affected by IL-1Ra treatment.

**Figure 5.**
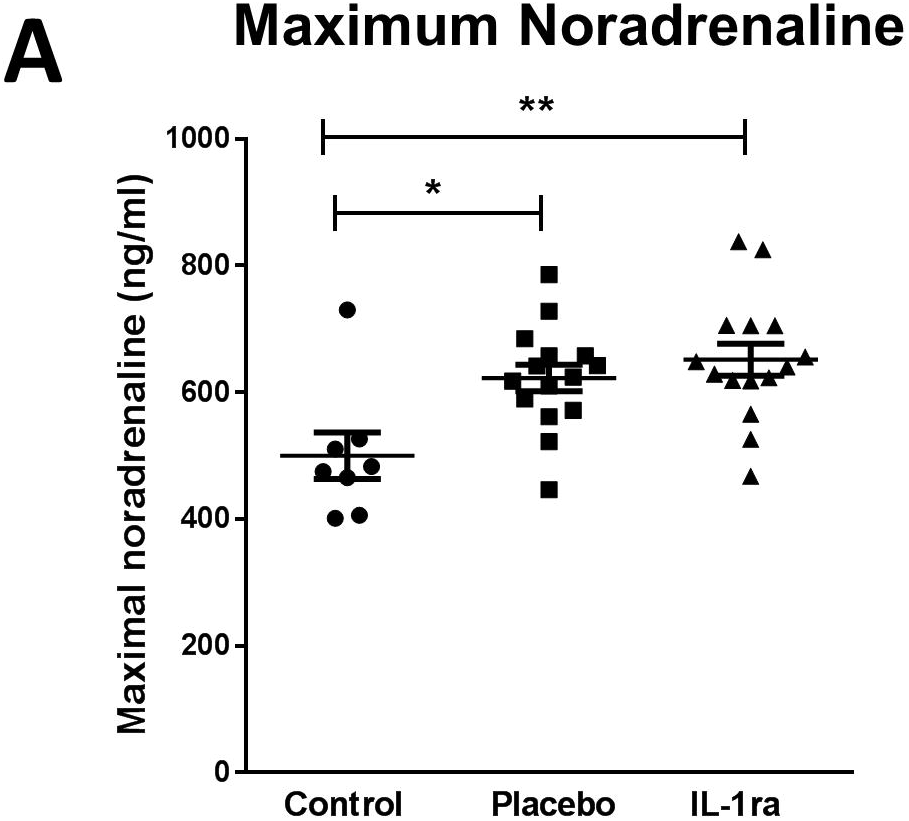
Plasma noradrenaline concentration is increased after stroke and is not affected by treatment with IL-1Ra. **(A)** Maximal noradrenaline concentration measured in the first 7 d after stroke was significantly higher in both placebo and IL-1Ra treated patients in comparison to controls. Data show mean ± SD *\*P<0.05; **P<0.01;* one-way ANOVA.

## DISCUSSION

The IL-1 family of cytokines play a critical role in host defence to pathogens by signalling to a variety of host cells to induce downstream effects including, but not limited to, pro-inflammatory cytokine and chemokine production, immune cell recruitment and upregulation of vascular adhesion molecules ^10, 11^. However, in conditions of sterile inflammation and tissue injury, such as stroke, these effects can aggravate primary tissue damage and impair injury repair mechanisms. Blocking IL-1 signalling has shown improved outcome in both experimental animal and patient stroke studies ^1, 4, 12^. However, the immunosuppressive effects of blocking IL-1 signalling after stroke may additionally inhibit systemic responses to infection, further increasing the risk of infection in patients who are already immune compromised ^13, 14^. Indeed, meta-analysis studies have shown an increased risk of serious infection in rheumatoid arthritis patients treated for prolonged periods with the IL-1 blocking drug anakinra ^13^. However as of yet this has not been observed in stroke patients potentially reflecting differences in the duration of treatment. No statistically significant differences in infection incidence were seen between IL-1Ra and placebo treated patients in this study with 5/17 IL-1Ra treated patients experiencing infection between admission and d7 and 4/17 infections in placebo treated patients. Consistent with this pattern we have shown here that relatively short duration of treatment with IL-1Ra after acute stroke did not further affect stroke-induced changes to circulating immunoglobulin, complement or noradrenaline concentrations and is therefore unlikely to further compromise immune defence against infection through reducing the availability of these antibacterial mediators.

IL-1 cytokine family members are reported to have variable effects on B cell antibody production. IL-1β was reported to be important for the rapid production of anti-bacterial IgM by innate-like B cells important for early containment of infection prior to the induction of adaptive immune responses ^7, 15^. This would suggest treatment with IL-1Ra after stroke could further compromise the early production of IgM in innate-like B cells which are already known to be reduced in number after stroke ^6^. However, this effect of IL-1Ra on IgM concentrations was not seen. We know that experimental stroke results in a significant loss of many populations of B cells and associated IgM ^6^, therefore it is possible that the effects of the stroke itself on B cells overwhelm any additional effects of cytokines that could moderately enhance or inhibit immunoglobulin production. Furthermore, we do not know if remaining B cells are functionally impaired and therefore able to respond to IL-1β signalling as they would under normal homeostatic conditions. We have previously reported that stroke is associated with reduced circulating IgM concentrations in comparison to non-stroke controls ^6^, an effect reproduced here. Further studies will be required to determine if IgM, or any of the mediators assessed in this study, would be useful as biomarkers to determine which patients are likely to develop infection after stroke.

We have shown, for the first time that circulating IgG1, IgG4 and IgA concentrations were reduced in the first 7 d after stroke in comparison to non-stroke controls. This is in agreement with previous data showing that pan-IgG concentrations were reduced in patients after stroke although subclasses of IgG were not assessed in that study and no reduction in IgA was found at the 7 d time point assessed ^16^. IgA is the most predominant immunoglobulin isotype at mucosal surfaces including the respiratory tract and is crucial for antibacterial protection at these sites ^17^. Given the early reduction of IgA in placebo-treated stroke patients, determining the effect of stroke on IgA-producing B cells at infection susceptible sites such as the lung mucosa could further elucidate if this has an important role in post-stroke infection susceptibility.

In contrast to the short half-life of IgA and IgM ^17–19^, the half-lives of IgG1 and IgG4 are reported to be 21 d and therefore an early reduction in IgG concentration is not compatible with a lack of *de novo* production after stroke due to loss of B cells ^20^. Previous studies have suggested that reduced total-IgG after stroke may be associated with increased loss or catabolism of IgG which could account for reductions in concentration occurring more rapidly than its natural half-life ^21^. An alternative explanation could be that reduced IgG concentration is indicative of vascular risk factors and inflammatory changes preceding stroke that are associated with stroke risk. However, control patients in this study were matched for risk factors including their degree of atherosclerosis and would be expected to show similar changes to stroke patients if these were associated with risk factors. Understanding the kinetics of individual immunoglobulin subset changes both preceding, and as a result of stroke, and their associations with post-stroke infections, could be invaluable in providing new therapeutic targets to reduce incidence of infection and improve outcome in patients.

The complement system has a crucial role in enhancing humoral immune defence and protecting from bacterial infection via interactions with both the innate and adaptive immune systems ^22^. As activation of complement is closely associated with efficient immunoglobulin-mediated clearance of pathogens, we determined if these pathways were compromised by stroke. We have assessed for the first time, individual concentrations of multiple complement components covering all pathways of complement activation after stroke. These exploratory data suggest there are no overall deficits in complement activation after stroke. Complement activation pathways converge at multiple points, however their initial activation mechanisms are distinct. The classical complement pathway is activated when IgM or IgG immune complexes bind to C1 (composed of C1q, C1r and C1s) ^22, 23^. Maximum circulating concentration of complement components associated with the classical and lectin pathways of activation, C1q, C2 and C4b and end stage mediators common to all pathways, C5 and C9 were increased in the first 7 d after stroke in comparison to non-stroke controls. As concentrations of MBL itself was not significantly altered by stroke, this suggests the classical complement pathway is specifically activated after stroke.

In contrast, the alternative pathway of complement activation is initiated by microbial cell surfaces and polysachharide antigen and results in a cascade that generates C3 ^22, 23^. Complement components that were significantly downregulated after stroke, C3b/ iC3b, C3, Factor H (fH) and Properdin, are more associated with the alternative pathway of complement activation, suggesting that the alternative pathway is suppressed. These data are in agreement with previous studies investigating systemic CRP, C3c and C4 complement concentrations in the serum of patients 24 h after ischemic stroke which concluded the classical pathway of complement activation was activated in the first 24 h after ischemic stroke whereas C3c, associated with the alternative pathway, was reduced ^24, 25^. The roles of individual pathways of complement activation in infection susceptibility after stroke remains to be determined but these data suggest overall deficits in complement concentration are unlikely to contribute to reduced antibody-mediated clearance of pathogens that may occur after stroke further supporting reduced circulating immunoglobulins as an important influence on infection susceptibility.

In this study, circulating noradrenaline concentrations measured in the first week after stroke were increased in comparison to non-stroke controls but were not influenced by treatment with IL-1Ra. This is in agreement with previous studies showing activation of the sympathetic nervous system in both stroke and subarachnoid haemorrhage patients that resulted in increased plasma noradrenaline concentrations that persisted up to 10 days ^26–28^. Our previous studies have shown that after experimental stroke, activation of the sympathetic nervous system and release of noradrenaline within the spleen is toxic to resident B cells and preventing noradrenaline signalling using the β-blocker propranolol prevented B cell and IgM loss and resulted in reduced infectious burden ^6^. The cytokine IL-1β is also increased in the spleen after stroke and is reported to activate peripheral nerves, including the splenic nerve, and increase production of splenic noradrenaline ^29, 30^. However blockade of IL-1β signalling did not alter circulating concentrations of noradrenaline after stroke.

In summary, we have shown that treatment with IL-1Ra after stroke does not affect circulating concentrations of immunoglobulins, complement components or noradrenaline and is therefore unlikely to further increase patient susceptibility to infection via pathways in which these mediators are key participants. This is in agreement with data from IL-1Ra Phase 2 trials in which treatment of stroke patients with IL-1Ra did not aggravate incidence of infection ^4, 31^. These data suggest that blocking IL-1 in a stroke context may not be concerning from the perspective of increasing infection risk in patients. Additionally, the reductions in circulating immunoglobulin concentrations detected after stroke in this study further support that antibody mediated immune defence may be an important therapeutic target to reduce the burden of infection after stroke.

## Acknowledgements

We thank Dr Hedley Emsley for recruitment of patients and data collection during the original stroke patient study, and for all the participating patients and controls for their participation and consent. We also thank Sharon Hulme for assistance with ethical applications and sample transfer. We thank Merck Millipore Corporation, Billerica, MA, USA for kind provision of the Milliplex^®^_MAP_ immunoglobulin isotyping and complement panel kits used in this study.

## Abbreviations

CRP: C-Reactive protein
IL-1Ra: IL-1 receptor antagonist
MBL: mannose-binding lectin
NIHSS: National Institute of Health Stroke Scale
SNS: sympathetic nervous system
WBC: White blood cell

## References

1. Sobowale OA, Parry-Jones AR, Smith CJ, Tyrrell PJ, Rothwell NJ, Allan SM. Interleukin-1 in Stroke. From Bench to Bedside 2016;47:2160–2167.

2. Touzani O, Boutin H, Chuquet J, Rothwell NJ. Potential mechanisms of interleukin-1 involvement in cerebral ischaemia. Journal of Neuroimmunology 1999;100:203–215.

3. Pradillo JM, Denes A, Greenhalgh AD, et al. Delayed Administration of Interleukin-1 Receptor Antagonist Reduces Ischemic Brain Damage and Inflammation in Comorbid Rats. Journal of Cerebral Blood Flow & Metabolism 2012;32:1810–1819.

4. Emsley HCA, Smith CJ, Georgiou RF, et al. A randomised phase II study of interleukin-1 receptor antagonist in acute stroke patients. Journal of Neurology, Neurosurgery & Psychiatry 2005;76:1366–1372.

5. Iadecola C, Anrather J. The immunology of stroke: from mechanisms to translation. Nat Med 2011;17:796–808.

6. McCulloch L, Smith CJ, McColl BW. Adrenergic-mediated loss of splenic marginal zone B cells contributes to infection susceptibility after stroke. Nature Communications 2017;8:15051.

7. del Barrio L, Sahoo M, Lantier L, Reynolds JM, Ceballos-Olvera I, Re F. Production of Anti-LPS IgM by B1a B Cells Depends on IL-1β and Is Protective against Lung Infection with Francisella tularensis LVS. PLOS Pathogens 2015;11:e1004706.

8. Martin F, Oliver AM, Kearney JF. Marginal Zone and B1 B Cells Unite in the Early Response against T-Independent Blood-Borne Particulate Antigens. Immunity 2001;14:617–629.

9. Baumgarth N, Herman OC, Jager GC, Brown L, Herzenberg LA, Herzenberg LA. Innate and acquired humoral immunities to influenza virus are mediated by distinct arms of the immune system. Proceedings of the National Academy of Sciences 1999;96:2250–2255.

10. Palomo J, Dietrich D, Martin P, Palmer G, Gabay C. The interleukin (IL)-1 cytokine family – Balance between agonists and antagonists in inflammatory diseases. Cytokine 2015;76:25–37.

11. Dinarello CA, Simon A, van der Meer JWM. Treating inflammation by blocking interleukin-1 in a broad spectrum of diseases. Nature Reviews Drug Discovery 2012;11:633.

12. Rothwell NJ. Interleukin-1 and neuronal injury: mechanisms, modification, and therapeutic potential. Brain, Behavior, and Immunity 2003;17:152–157.

13. Salliot C, Dougados M, Gossec L. Risk of serious infections during rituximab, abatacept and anakinra treatments for rheumatoid arthritis: meta-analyses of randomised placebo-controlled trials. Annals of the Rheumatic Diseases 2009;68:25–32.

14. Westendorp W, Nederkoorn P, Vermeij J-D, Dijkgraaf M, van de Beek D. Post-stroke infection: A systematic review and meta-analysis. BMC Neurology 2011;11:110.

15. Zouali M, Richard Y. Marginal zone B-cells, a gatekeeper of innate immunity. Frontiers in Immunology 2011;2.

16. Liesz A, Roth S, Zorn M, Sun L, Hofmann K, Veltkamp R. Acquired Immunoglobulin G deficiency in stroke patients and experimental brain ischemia. Experimental Neurology 2015;271:46–52.

17. Mkaddem SB, Christou I, Rossato E, Berthelot L, Lehuen A, Monteiro RC. IgA, IgA Receptors, and Their Anti-inflammatory Properties. In: Daeron M, Nimmerjahn F, eds. Fc Receptors. Cham: Springer International Publishing, 2014: 221–235.

18. Fahey JL, Sell S. THE IMMUNOGLOBULINS OF MICE: V. THE METABOLIC (CATABOLIC) PROPERTIES OF FIVE IMMUNOGLOBULIN CLASSES. The Journal of Experimental Medicine 1965;122:41–58.

19. Sigounas G, Harindranath N, Donadel G, Notkins AL. Half-life of polyreactive antibodies. Journal of Clinical Immunology 1994;14:134–140.

20. Morell A, Terry WD, Waldmann TA. IgG subclasses: Physical properties, genetics and biological functions. J Clin Invest 1970;1970:673–680.

21. Liesz A, Dalpke A, Mracsko E, et al. DAMP Signaling is a Key Pathway Inducing Immune Modulation after Brain Injury. The Journal of Neuroscience 2015;35:583–598.

22. Kemper C, Atkinson JP. T-cell regulation: with complements from innate immunity. Nature Reviews Immunology 2006;7:9.

23. Holers MV. Complement and Its Receptors: New Insights into Human Disease. Annual Review of Immunology 2014;32:433–459.

24. Pedersen ED, Waje-Andreassen U, Vedeler CA, Aamodt G, Mollnes TE. Systemic complement activation following human acute ischaemic stroke. Clinical & Experimental Immunology 2004;137:117–122.

25. Di Napoli M. Systemic Complement Activation in Ischemic Stroke. Stroke 2001;32:1443–1448.

26. Naredi S, Lambert G, Edén E, et al. Increased sympathetic nervous activity in patients with non-traumatic subarachnoid hemorrhage. Stroke 2018;31:901–906.

27. Urra X, Cervera Á, Obach V, Climent N, Planas AM, Chamorro Á. Monocytes Are Major Players in the Prognosis and Risk of Infection After Acute Stroke. Stroke 2009;40:1262–1268.

28. Chamorro A, Amaro S, Vargas M, et al. Catecholamines, infection, and death in acute ischemic stroke. J Neurol Sci 2007;252.

29. Schwarting S, Litwak S, Hao W, Bähr M, Weise J, Neumann H. Hematopoietic Stem Cells Reduce Postischemic Inflammation and Ameliorate Ischemic Brain Injury. Stroke 2008;39:2867–2875.

30. Niijima A, Hori T, Aou S, Oomura Y. The effects of interleukin-1β on the activity of adrenal, splenic and renal sympathetic nerves in the rat. Journal of the Autonomic Nervous System 1991;36:183–192.

31. Smith CJ, Hulme S, Vail A, et al. SCIL-STROKE (Subcutaneous Interleukin-1 Receptor Antagonist in Ischemic Stroke). A Randomized Controlled Phase 2 Trial 2018;49:1210–1216.

