## Supplementary Figures 1-4 for "Interleukin-1 receptor antagonist treatment in acute ischaemic stroke does not alter systemic markers of anti-microbial defence"

### Slide 1
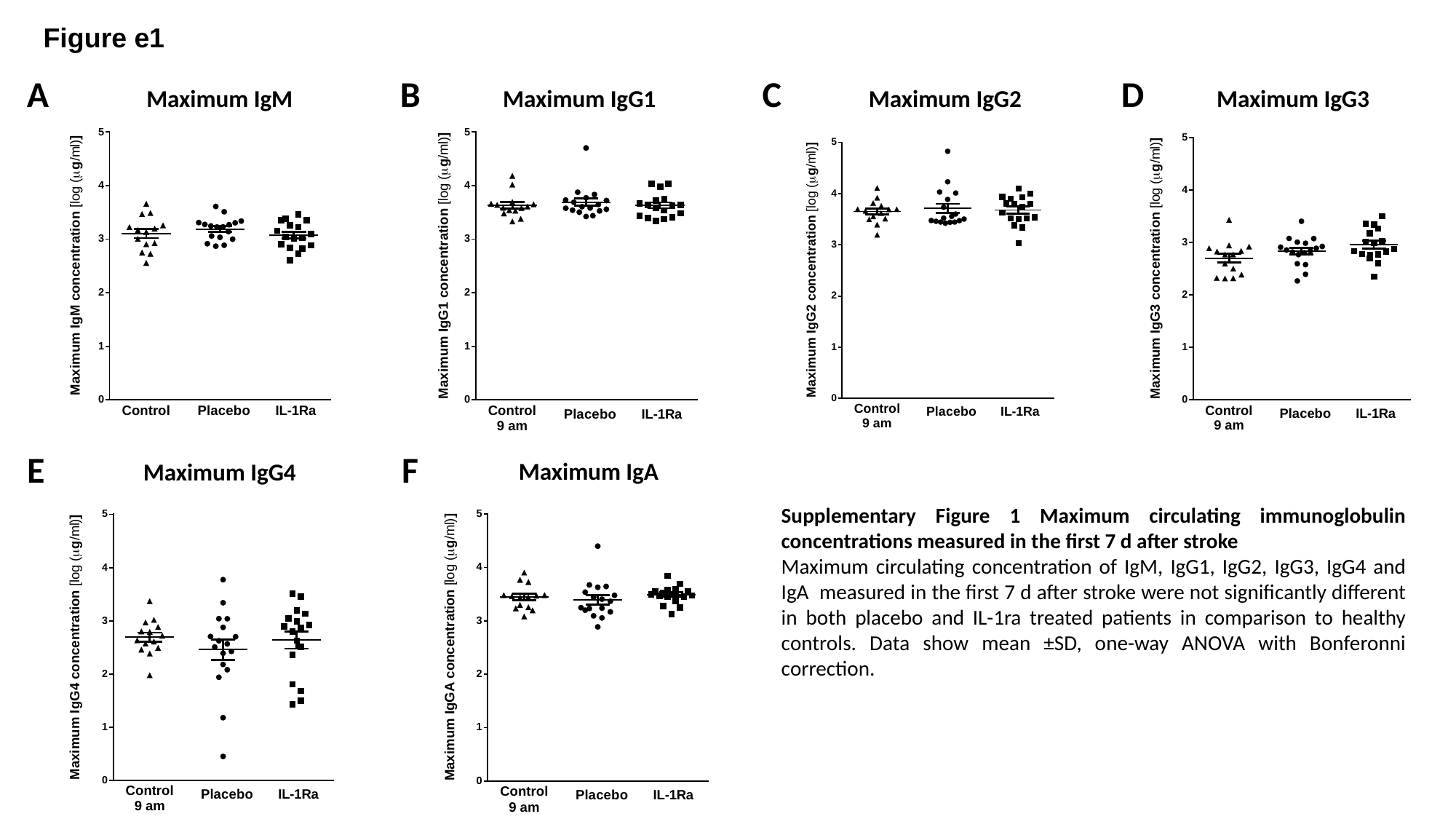

Figure e1
A
B
C
D
Maximum IgM
Maximum IgG1
Maximum IgG2
Maximum IgG3
E
F
Maximum IgA
Maximum IgG4
Supplementary Figure 1 Maximum circulating immunoglobulin concentrations measured in the first 7 d after stroke
Maximum circulating concentration of IgM, IgG1, IgG2, IgG3, IgG4 and IgA measured in the first 7 d after stroke were not significantly different in both placebo and IL-1ra treated patients in comparison to healthy controls. Data show mean ±SD, one-way ANOVA with Bonferonni correction.

### Slide 2
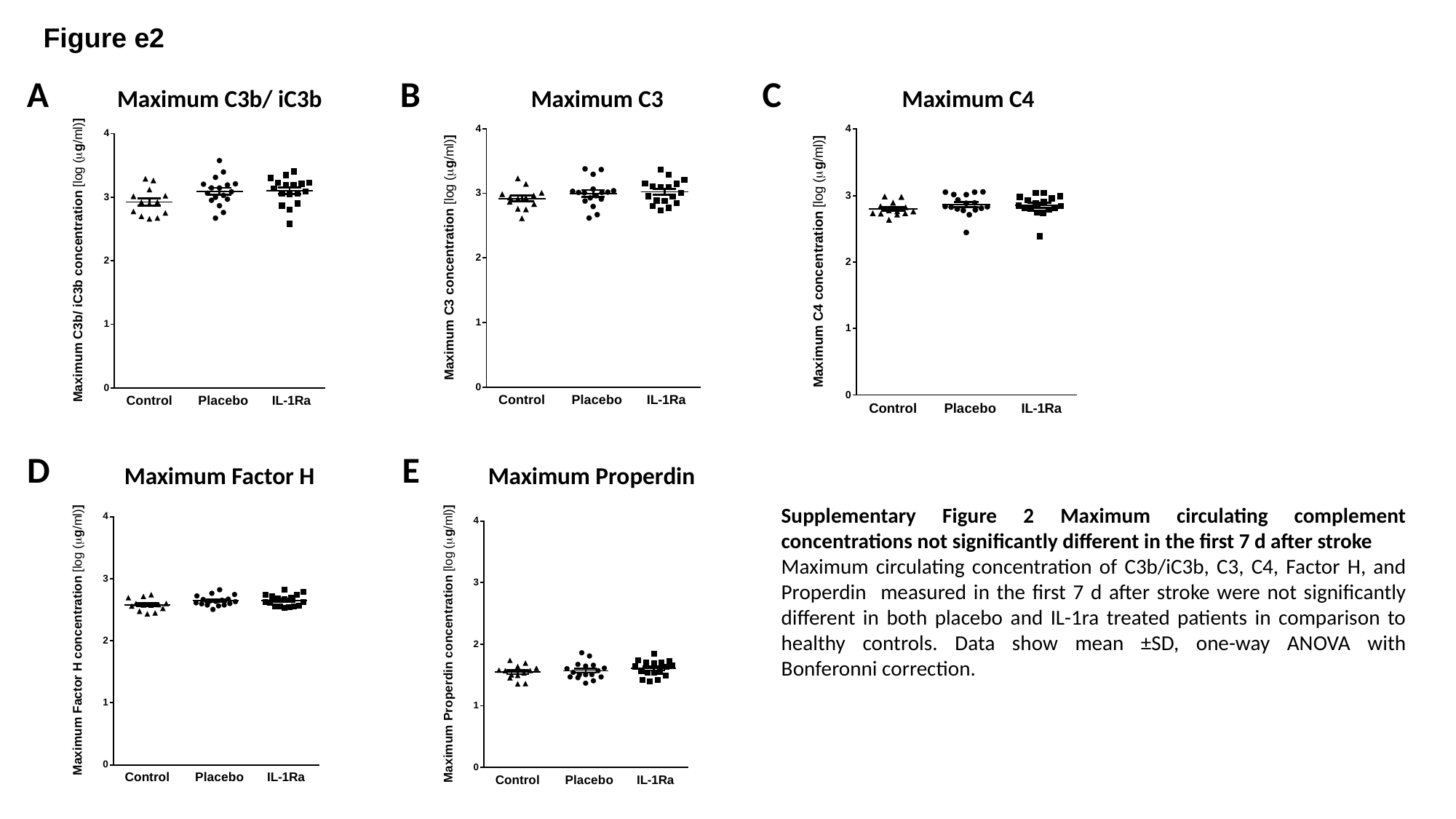

Figure e2
A
B
C
Maximum C3b/ iC3b
Maximum C3
Maximum C4
D
E
Maximum Factor H
Maximum Properdin
Supplementary Figure 2 Maximum circulating complement concentrations not significantly different in the first 7 d after stroke
Maximum circulating concentration of C3b/iC3b, C3, C4, Factor H, and Properdin measured in the first 7 d after stroke were not significantly different in both placebo and IL-1ra treated patients in comparison to healthy controls. Data show mean ±SD, one-way ANOVA with Bonferonni correction.

### Slide 3
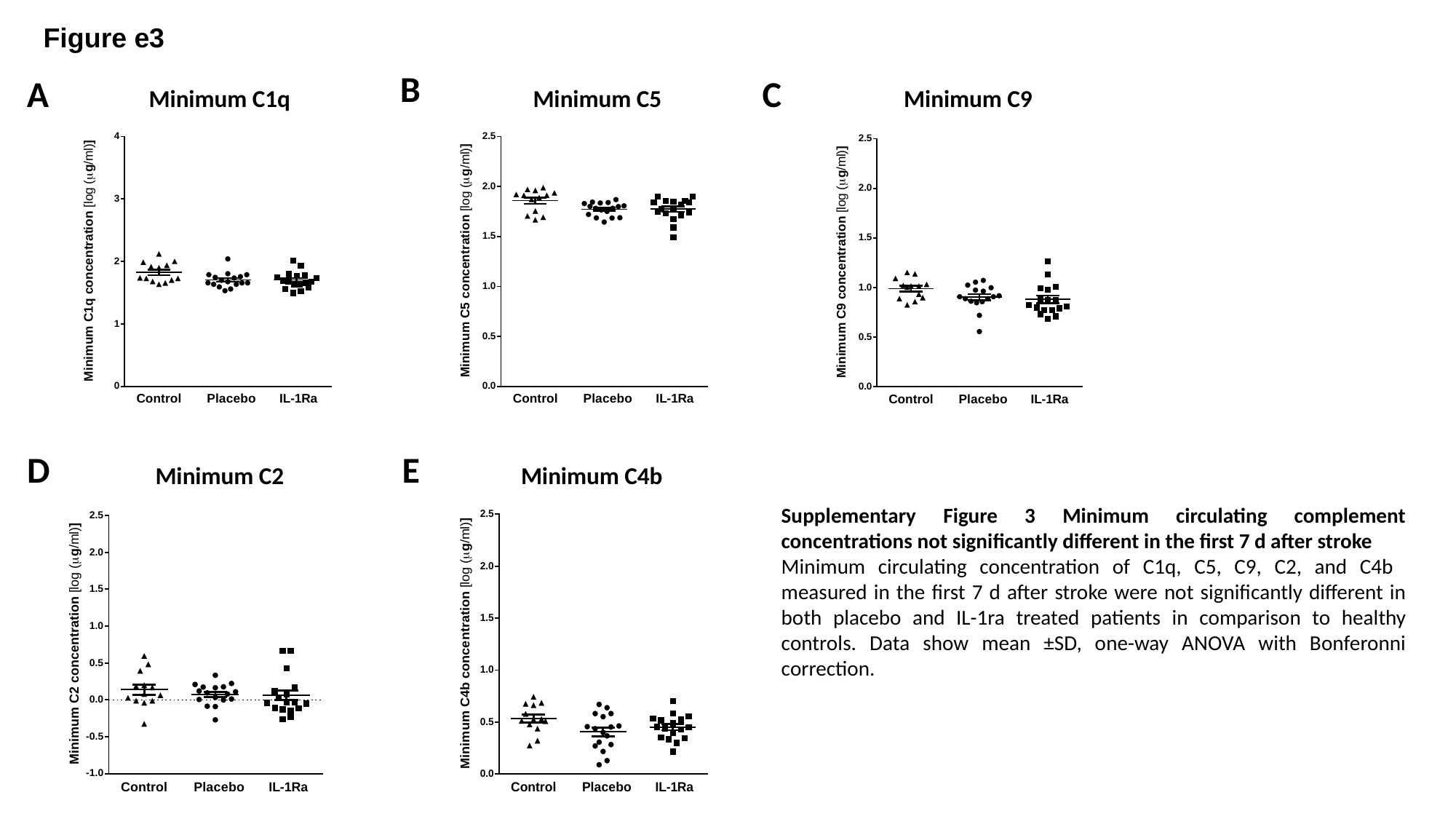

Figure e3
B
A
C
Minimum C1q
Minimum C5
Minimum C9
D
E
Minimum C2
Minimum C4b
Supplementary Figure 3 Minimum circulating complement concentrations not significantly different in the first 7 d after stroke
Minimum circulating concentration of C1q, C5, C9, C2, and C4b measured in the first 7 d after stroke were not significantly different in both placebo and IL-1ra treated patients in comparison to healthy controls. Data show mean ±SD, one-way ANOVA with Bonferonni correction.

### Slide 4
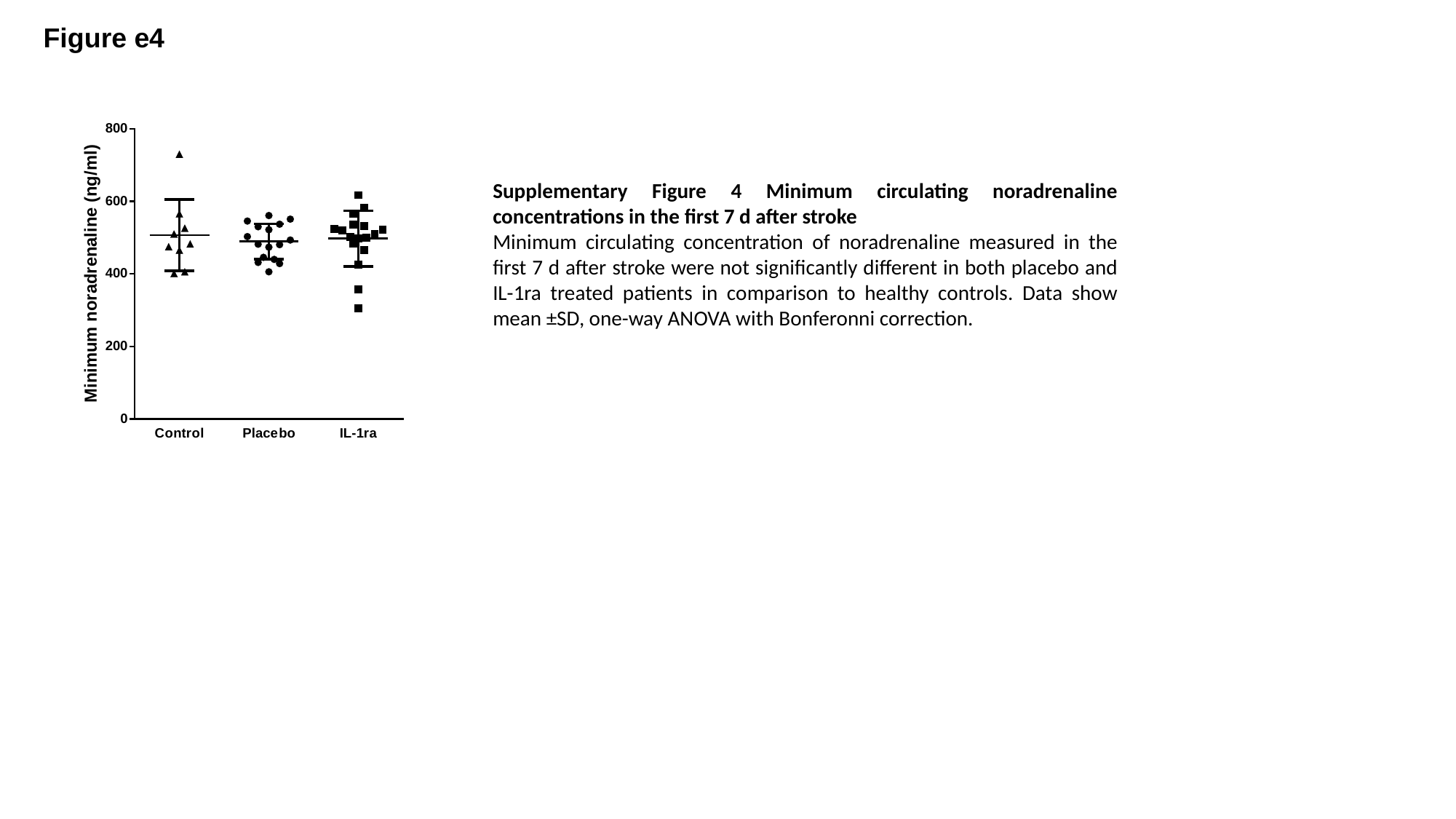

Figure e4
Supplementary Figure 4 Minimum circulating noradrenaline concentrations in the first 7 d after stroke
Minimum circulating concentration of noradrenaline measured in the first 7 d after stroke were not significantly different in both placebo and IL-1ra treated patients in comparison to healthy controls. Data show mean ±SD, one-way ANOVA with Bonferonni correction.

### Slide 5
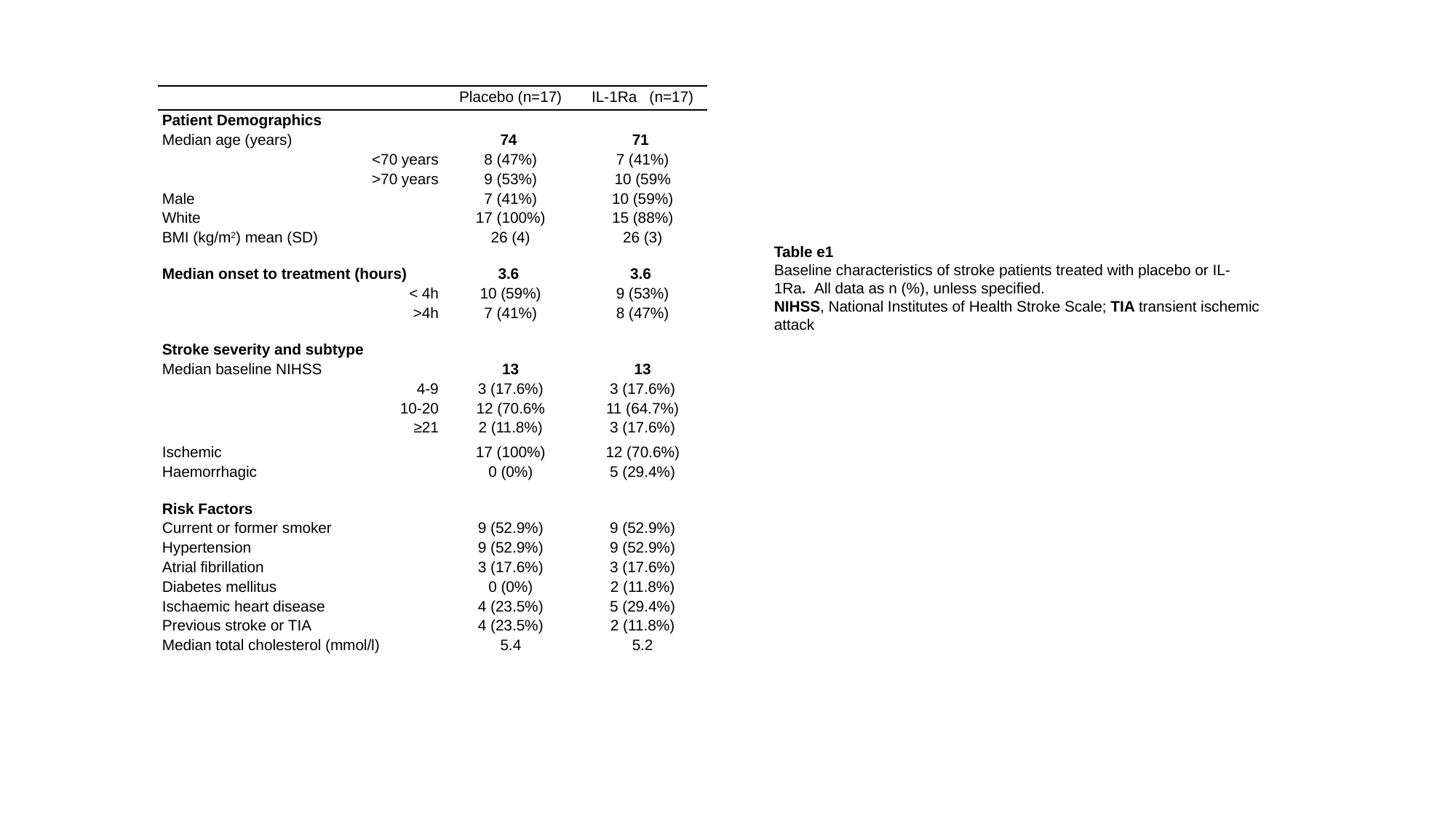

| | | |
| --- | --- | --- |
| | Placebo (n=17) | IL-1Ra (n=17) |
| Patient Demographics | | |
| Median age (years) | 74 | 71 |
| <70 years | 8 (47%) | 7 (41%) |
| >70 years | 9 (53%) | 10 (59% |
| Male | 7 (41%) | 10 (59%) |
| White | 17 (100%) | 15 (88%) |
| BMI (kg/m^2^) mean (SD) | 26 (4) | 26 (3) |
| Median onset to treatment (hours) | 3.6 | 3.6 |
| < 4h | 10 (59%) | 9 (53%) |
| >4h | 7 (41%) | 8 (47%) |
| Stroke severity and subtype | | |
| Median baseline NIHSS | 13 | 13 |
| 4-9 | 3 (17.6%) | 3 (17.6%) |
| 10-20 | 12 (70.6% | 11 (64.7%) |
| ≥21 | 2 (11.8%) | 3 (17.6%) |
| Ischemic | 17 (100%) | 12 (70.6%) |
| Haemorrhagic | 0 (0%) | 5 (29.4%) |
| Risk Factors | | |
| Current or former smoker | 9 (52.9%) | 9 (52.9%) |
| Hypertension | 9 (52.9%) | 9 (52.9%) |
| Atrial fibrillation | 3 (17.6%) | 3 (17.6%) |
| Diabetes mellitus | 0 (0%) | 2 (11.8%) |
| Ischaemic heart disease | 4 (23.5%) | 5 (29.4%) |
| Previous stroke or TIA | 4 (23.5%) | 2 (11.8%) |
| Median total cholesterol (mmol/l) | 5.4 | 5.2 |
Table e1
Baseline characteristics of stroke patients treated with placebo or IL-1Ra. All data as n (%), unless specified.
NIHSS, National Institutes of Health Stroke Scale; TIA transient ischemic attack
